# Assessing rapid adaptation through epigenetic inheritance: a new experimental approach

**DOI:** 10.1101/2023.10.12.562085

**Authors:** Alexandra Chávez, Meret Huber

## Abstract

1. Epigenetic inheritance is hypothesized to mediate rapid adaptation to stresses via two fundamentally different routes: first, through spontaneous epimutations that arise in a largely stochastic manner in the presence or absence of stress; if these spontaneous epimutations are heritable and beneficial, they may be selected upon (“stochastic route”); and second, through environment-induced epialleles that arise uniformly among individuals; if heritable, these epialleles may lead to stress adaptation even in the absence of selection (“deterministic route”). Testing and teasing apart these two routes is challenging, largely because a suitable experimental approach is lacking.
2. Here, we propose an experimental approach that allows to simultaneously assess the contribution of the stochastic and deterministic route. The essence of the approach is to manipulate the efficacy of selection through the population size and thereby to test whether selection is required for adaptation (stochastic route). To this end, genetically uniform populations are grown under different environments across multiple generations (“pre-treatment”) at two different population sizes: in large populations, in which selection is effective; and in small populations, in which drift overcomes the effect of selection. If the deterministic route contributes to adaptation, variation in fitness, phenotypes or epigenetic marks will arise between the small populations of the different pre-treatments. If the stochastic route contributes to adaptation, variation will arise between the small and large population within each pre-treatment. As a proof-of-principle, we tested whether small and large monoclonal populations of the aquatic duckweed *Spirodela polyrhiza* may adapt to copper excess outdoors.
3. After five to seven generations of pre-treatment and a subsequent multi-generational growth under control conditions, large populations outperformed small populations under copper excess. Furthermore, small populations pre-treated with copper excess tended to have higher fitness under copper excess than small populations pre-treated under control conditions. These data suggest that both the stochastic and deterministic route may alter plant fitness under recurring stress.
4. The proposed approach will allow to experimentally evaluate whether species may adapt to stresses through either stochastic and deterministic epigenetic changes, which is fundamental to understand whether and how epigenetic inheritance may lead to rapid stress adaptation.

## INTRODUCTION

The hallmark of evolutionary biology states that species adapt to stresses through selection of phenotypic variants that are arise from DNA sequence variation. Yet, increasing evidence shows that DNA sequence variation is not the sole source of heritable phenotypic variation: for instance, flowers of natural toadflax mutants develop radial instead of bilateral symmetry because a floral symmetry gene became hypermethylated (Cubas et al., 1999). Similarly, homeotic floral phenotypes in the oil palm arose during tissue culture because an homeotic gene is alternatively spliced upon spontaneous hypomethylation of an intron located transposon (Ong-Abdullah et al., 2015). Furthermore, fruit ripening in a spontenous tomato mutant is hampered because a gene of the SBP-box family of trascription factors became hypermerthylated and was thereby silenced (Manning et al., 2006). These examples highlight that mechanisms other than DNA sequence variation may lead to heritable, variable and fitness-relevant phenotypes, and as such they open up an exciting question: which role do mechanisms that generate heritable phenotypic variation in the absence of DNA sequence variation play in the adaptation to environmental stresses? Considering the on-going loss of intra-specific genetic diversity and the rapid pace of environmental change (Des Roches et al., 2021; Sage, 2020), answering this question is becoming an increasingly relevant.

Heritable phenotypic variation that is not caused by DNA sequence changes may arise through so-called “non-genetic” or “epigenetic” inheritance (Bonduriansky & Day, 2009; O’Dea et al., 2016). Epigenetic inheritance often refers to mechanisms that alter gene expression across generations through genome-associated mechanisms such as DNA methylation, histone modifications and small RNAs, whereas non-genetic inheritance includes any other mechanism such as the vertical transfer of microbes, nutrients or hormones (Bonduriansky & Day, 2009). In this manuscript, we use the term “epigenetic” to refer to both epigenetic and other non-genetic mechanisms in order to improve readibily.

Many epigenetic marks share two features: first, epigenetic marks may change spontaneously over generations in a stochastic manner (Becker et al., 2011; Schmitz et al., 2011), and second, epigenetic marks may be induced by the environment and thereby lead to predictable phenotypic variation (Wibowo et al., 2016; Zhang et al., 2018). Consequently, epigenetic inheritance may lead to stress adaptation through two fundamentally different routes: the “stochastic” and the “deterministic” route (Fig. 1a) (Baugh & Day, 2020).

**Figure 1.**
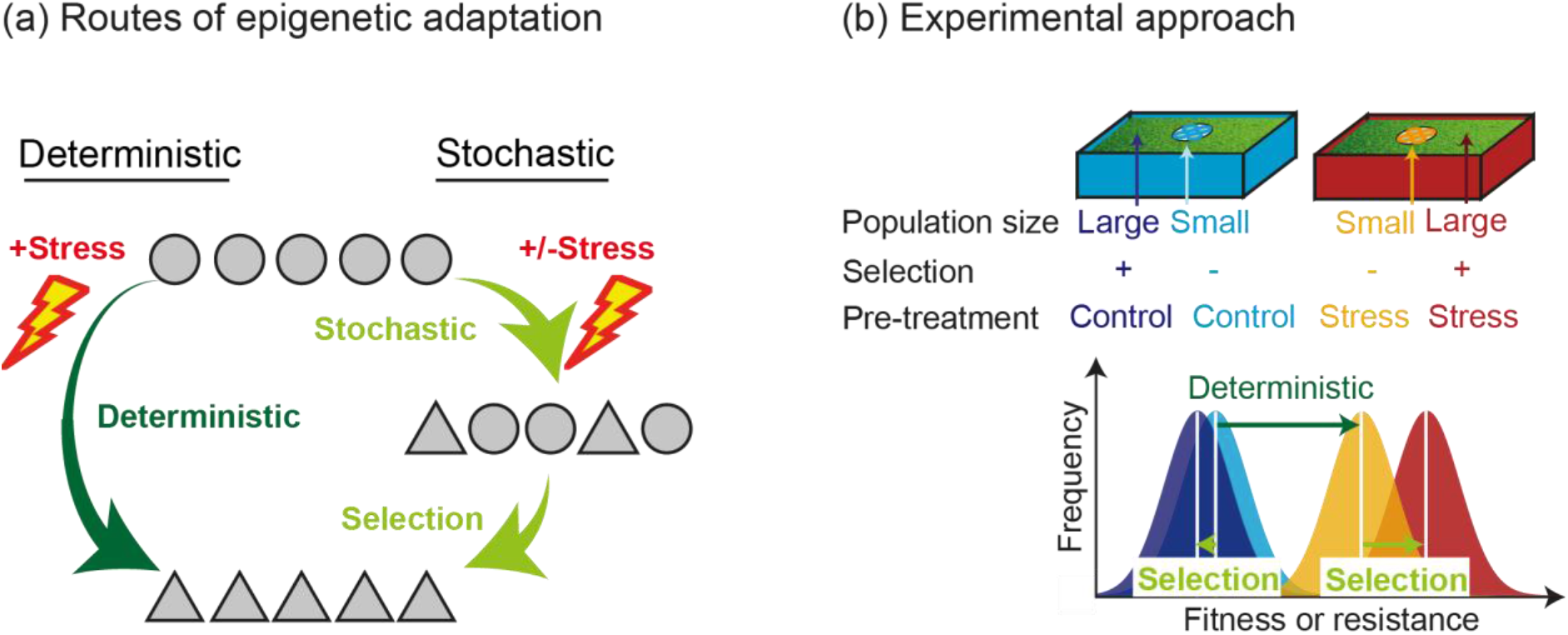
The two routes of rapid adaptation through epigenetic inheritance. (a) Epigenetic inheritance may facilitate adaptation either via deterministic epialleles (deterministic route) or selection of stochastic epigenetic variants (stochastic route). Each circle and triangle indicate the DNA methylation level of a specific locus of different individuals. (b) To differentiate among the deterministic and stochastic route, organisms are grown in two different environments (pre-treatments) in large and small populations; after a multi-generational recovery phase in a shared control environment, fitness, phenotypes and epigenetic marks in the presence and absence of recurring stress are compared between populations sizes and pre-treatments.

The stochastic route is comparable to the adaptive process that is based on DNA sequence variation. Here, spontaneous changes (“epimutations”) arise either in the presence or absence of stress in a largely random manner. For instance, in *Arabidopsis thaliana*, variations in CG and to a much lower extent also CHG and CHH methylation (H=A, T, C) steadily accumulate across generations (Becker et al., 2011; Denkena et al., 2021; Schmitz et al., 2011), and both the frequency and the spectrum of these spontaneous epimutations may be altered by stress (Jiang et al., 2014; Johannes & Schmitz, 2019). In plants, these spontaneous epimutations are semi-stable, meaning that many epimutations are inherited across several generations but also frequently reverse (Johannes & Schmitz, 2019). Spontaneous epimutations increase the phenotypic diversity in genetically uniform individuals upon which natural selection can act.

The deterministic route, in contrast, is not comparable to the adaptive process that is based on DNA sequence variation. In the deterministic route, also described as transgenerational plasticity, reviewed in Anastasiadi et al. (2021), environmental stresses induce epigenetic variation and thereby lead to predictable epigenetic variations that are shared among individuals (“deterministic epialleles”). For instance, in *A. thaliana,* environmental stresses may remodel CHG and CHH methylation in specific repeat sequences, which is associated with altered expression of stress-related genes nearby (Annacondia et al., 2021; Luna et al., 2012; Wibowo et al., 2016). Such stress-induced epialleles are usually considered to have limited heritability, as they often vanish after maximally three offspring generation (Luna et al., 2012; Wibowo et al., 2016), and thus may occur due to direct stress exposure during the early development of the offspring rather than being truly inherited. Recent evidence, however, indicates that deterministic epialleles may be substantially more stable than anticipated: in the duckweed *Lemna minor*, heat stress remodelled CHG-methylation, and part of this variation was still observed after at least three clonal generations under control conditions (Van Antro et al., 2023). Similarly, in the closely related duckweed *S. polyrhiza*, copper excess induced variation in phenotypes and fitness that persisted for up to 10 clonal generations under control conditions (Huber et al., 2021). If deterministic epialleles are heritable across multiple generations, as these studies suggest, they may lead to rapid stress adaption even in the absence of selection.

## METHODS

### The problem: teasing apart the stochastic and deterministic routes

While the stochastic route relies on random variation being selected upon, the deterministic route may allow populations to adapt in the absence of selection. Does epigenetic inheritance thereby lead to a fundamentally different adaptive process in which adaptation is, to a certain extent, predictable and deterministic, or does adaptation through epigenetic inheritance proceeds similar to genetic adaptation through selection of phenotypic variants that are generated through random epimutations? Answering this question is fundamental to understand whether and how epigenetic inheritance may lead to rapid adaption. Yet, assessing and teasing apart the stochastic and deterministic route is experimentally challenging.

Experiments that aim to test the deterministic route are relatively common, e.g. Baugh & Day (2020), Huber et al. (2021), Luna et al. (2012), Wibowo et al. (2016). Usually, genetically uniform organisms are grown for multiple generations under different environments (“pre-treatment”) as single descendants - thereby confounding genetic effects as well as selection of beneficial (epi-)genetic variants are almost completely avoided (Baugh & Day, 2020). Subsequently, individuals are grown for at least three generations under control conditions to avoid that the organisms were directly exposed to stress during their early development (Heard & Martienssen, 2014). Subsequently, fitness and epigenetic variation between the pre-treatments are assessed. Ideally this experiment is accompanied with genetic or chemical manipulation of the epigenetic machinery and/or genes that are transgenerationally regulated to test the molecular mechanisms. While this approach is powerful to test the role of deterministic epialleles in stress adaptation, it does not allow to assess whether selection of stochastic variants may lead to stress adaptation.

Experiments that test whether stochastic epigenetic variants may mediate rapid stress adaption through natural selection are rather rare, but see Heckwolf et al. (2020); Huber et al. (2021), Kronholm et al. (2017), Schmid et al. (2018), likely because of the experimental challenges. The most common approach is to grow large, genetically uniform populations for many generations under different environments (“pre-treatment”), subsequently grow individuals for a few generations under a shared control environment to erase environment-induced effects, and then compare fitness and epigenetic variation between the populations of the different pre-treatments. The limitation of this approach is that during the pre-treatment, populations simultaneously experience both selection of stochastic variants as well as the induction of deterministic epialleles – and if the deterministic epialleles are heritable for multiple generations, and show unexpected temporal inheritance patterns (Bell & Hellmann, 2019; Huber et al., 2021) – variation between the populations could be either due to selection of stochastic variants or heritable deterministic epialleles. To assess whether stochastic epigenetic variation may mediate rapid adaption through natural selection, we thus need an approach in which selection of stochastic variants can take place while teasing apart the contribution of deterministic epialleles.

### The solution: manipulating the efficacy of selection through the population size

Here, we propose an experimental approach that unifies both of the above-mentioned setups and thereby allows to simultaneously assess whether selection of stochastic variants and/or the formation of deterministic epialleles mediate rapid stress adaptation. One of the key differences between the stochastic and deterministic route is whether selection is needed – and as such, to differentiate among these two routes, one should manipulate the efficacy of selection. This can be achieved through the population size, as selection is most effective in large populations (McDonald, 2019; Wahl et al., 2002). We therefore propose to subject populations for several generations to different environments (“pre-treatments”) at two different population sizes: on the one hand, in population sizes sufficiently large that selection is effective. These populations may undergo both the stochastic and deterministic route. On the other hand, in population sizes sufficiently small that selection is not effective, i.e. that drift overcomes the effect of selection. This is achieved once the effective population size N_e_ < 1/s (s = selection coefficient) (McDonald, 2019; Nielsen & Slatkin, 2013); the smallest possible effective population size is achieved in single descendant lineages. Thus, the small population will evolve almost exclusively through drift, and thereby only undergo the deterministic route. Importantly, the large and small populations must be grown side by side in the very same environment to ensure that the deterministic route is equally induced in the small and large populations.

After several generations of pre-treatment, one can assess whether populations adapted to stress through the stochastic and/or deterministic route. To this end, individuals of the small and large populations of the different pre-treatments are grown in a shared control environment for at least three generations to ensure that any variation is truly heritable. Subsequently, phenotypes and fitness under the different environments are compared. This allows differentiating whether selection of epigenetic variants and/or deterministic epialleles lead to rapid adaptation: if deterministic epialleles confer resistance, variation in resistance and epigenetic variation will arise between the small populations of the different pre-treatments. If selection of epigenetic variants contributes to resistance, variation in resistance and in epigenetic variation will establish between the small and large population within each pre-treatment (Fig. 1b).

### The requirements: is my experimental system suitable for the approach?

While the proposed experiment is in principle applicable to species across the tree of life, the species under investigation should have following characteristics:

1. *Rapid reproduction and small body size.* As with all species used for experimental evolution, rapid reproduction and small body size are highly desirable.
2. *Low genetic variation.* Teasing apart genetic from epigenetic effects is notoriously difficult, see discussion below. We thus suggest minimizing genetic variation by using clonally reproducing organisms or organisms that are highly inbred; ideally, genetic mutation rates should be small to reduce genetic variation that arise during the experiment.
3. *Accurate assessments of fitness and epigenetic variation*. Measuring Darwinian fitness is critical when assessing adaptation – thus, direct fitness assessments (e.g. number of offspring) is preferred over indirect assessments such as phenotypes or altered performance of interacting species; such assessments are nevertheless valuable. Furthermore, the ability to measure epigenetic variation (e.g. DNA methylation, histone modification, small RNAs) in a largely unbiased manner and to link these variations to specific genes is highly desirable. Although these analyses often require a reference genome, new developments in high-throughput sequencing will likely overcome some of the limitations of non-model species that do not have a high-quality reference genome or whose genome is prohibitively large (Amarasinghe et al., 2020; Gawehns et al., 2022).

### Proof-of-concept: applying the method to assess rapid adaptation in monoclonal duckweed populations

A species that is ideal to test adaptation through the deterministic or stochastic route is the giant duckweed, *Spirodela polyrhiza,* one of the smallest and fastest growing flowering plants. *Spirodela polyrhiza* reproduces almost exclusively asexually by budding with a maximal doubling time of two to three days (Landolt, 1957; Ziegler et al., 2015). The plant consists of a flat, thallus-like shoot, the so-called frond, which allows to adequately estimate plant fitness by estimating growth rates based on surface area. The species has a very low mutation rate with approximately one point mutation every eighty generations (Xu et al., 2019). In nature, *S. polyrhiza* is exposed to diverse environmental stresses including copper excess, which is a common pollutant in aquatic ecosystems (Greger, 1999; Li & Xiong, 2004). To test whether *S. polyrhiza* acquires stress resistance through the stochastic or deterministic route, we grew monoclonal *S. polyrhiza* populations in the presence and absence of copper excess (“pre-treatment”) in both small and large populations (“selection regimen”) for approximately seven generations in ponds outdoors and assessed plant resistance to copper excess after five generations of growth in a shared control environment. The small populations consisted of single descendant lineages – the smallest possible population size – which were floating freely inside the large populations (Fig 2A).

**Figure 2.**
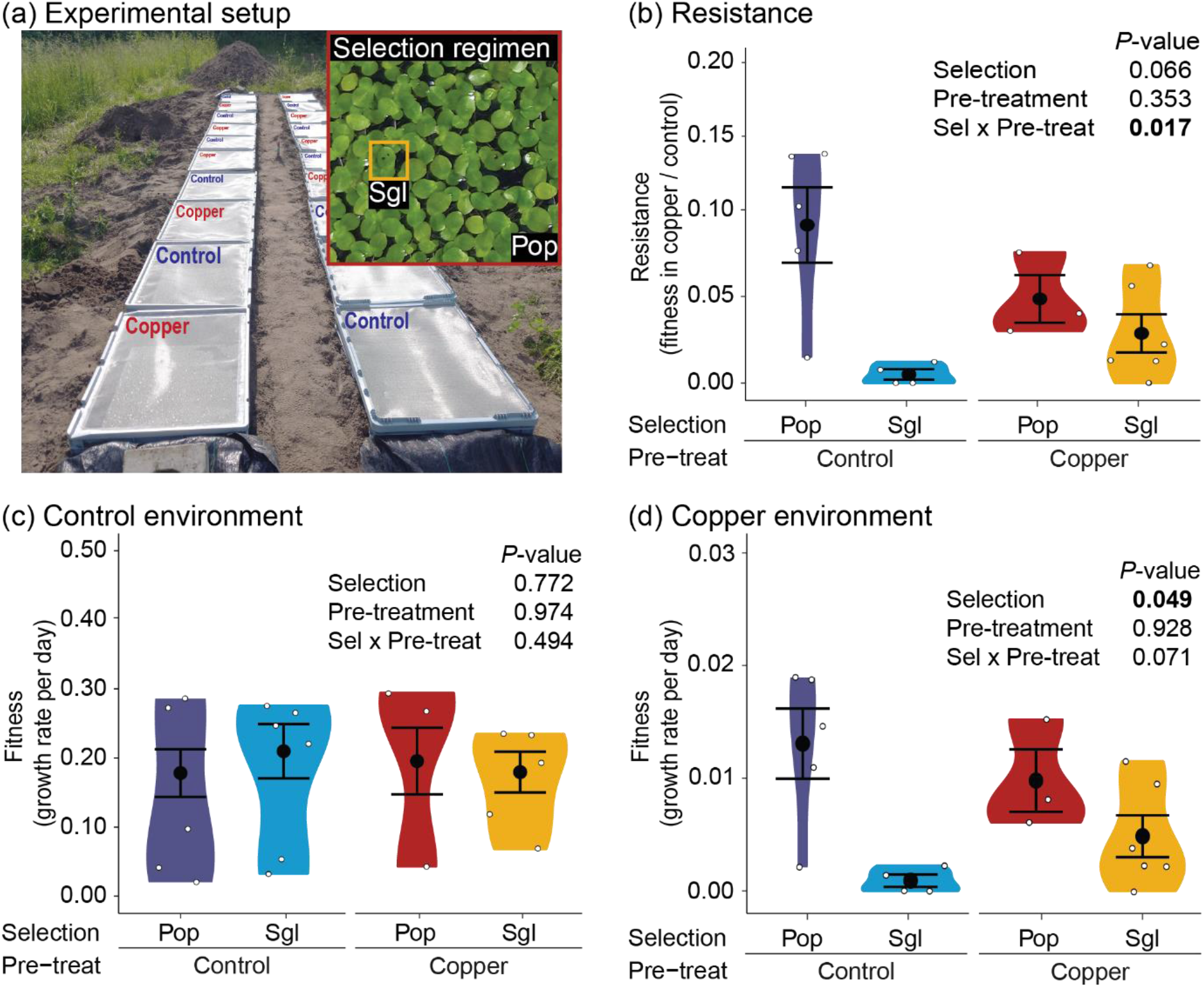
Testing the experimental approach by growing large and small monoclonal populations of Spirodela polyrhiza in the presence and absence of copper excess in outdoor mesocosms. (a) Overview of the experimental setup. (b) Plant resistance to copper excess (growth rates under copper excess compared to the group’s mean growth rate under control conditions) depended on the population size and the pre-treatment. (c) Growth rates under control environment was not affected by the population size and the pre-treatment environment. (d) Growth rates under copper excess were affected by the population size and the pre-treatment. For b-c: P-values refer to linear mixed effects models. The black circles show the means, and the error bars the standard errors. N = 3-6. Pre-treat = pre-treatment; Sel = selection; Pop = large population; Sgl = single descendants.

On May 2021, we set up 20 ponds (80 x 60 x 32cm, AuerPackaging, Amerang, Germany) inside an open field site in Münster, Germany (51°57’54.0’N, 7°36’22.4’E). Ponds were buried to the rim with soil and filled with 10 L of pond soil (Floragard, Oldenburg, Germany), 40 mL of organic fruit and vegetable fertilizer (organic NPK 3,1+0,5+4,1; COMPO, Münster, Germany) and tap water with a total volume of 100 L. The boxes had a draining scape and lids consisting of a stainless-steel mesh (mesh opening 0.63 mm, wire diameter, 0.224 mm; Haver&Boeker, Oelde, Germany). On May 26th, each box received around 3000 duckweed fronds of the genotype 7498 that were pre-cultivated inside 10% N-medium (Appenroth et al., 1996) for one month in indoor growth cabinets (GroBank, CLF PlantClimatics, Wertingen, Germany) operating at 26°C, 135 µmol photons m^-2^ s^-1^ supplied through LEDs with 16:8 h light:dark cycles. After one week of outdoor acclimation, we added 20 µM CuSO_4_ to half of the ponds (n=10) to initiate the stress pre-treatment (Fig. 2a). Across the entire experiments, we measured copper concentration weekly and after each rainfall; if needed, CuSO_4_ was added to maintain 20 µM CuSO_4._ After adding CuSO_4_ for the first time, we randomly chose four plants per pond as the founders of the single descendant lineages. To trace the single descendant lineages, we marked the plants with a dot using a permanent marker (Stabilo OHPen universal) (Fig. 2a). The remaining plants reproduced freely and covered the surface after 12 days, which corresponds to four to five generations of growth and a maximal population size of 25,000 individuals. To avoid crowding, we randomly removed 30% of the plants. Subsequently, we also marked four plants from the populations and followed them for two generations as single descendant lineages to equalize the marking treatment and the plant age between the single descendants and large populations. To grow the plants from the different pre-treatments and selection regimens in a uniform environment (“recovery phase”), we moved the third marked generation of the population and the sixth generation of the single descendants (approx. 24 days of pre-treatment) into separate cavities of polyvinyl chloride disks (7 cm diameter) that were floating inside the control ponds. Due to algae bloom, these plants did not reproduce any offspring. We thus transferred the very same plants back to the initial indoor conditions after 38 days of growth outdoors. To establish a shared environment among plants from the different pre-treatments and selection regimens, we placed the plants from the single descendant lineages and the large populations from each control and copper pre-treatment pond pair together inside a 250mL polypropylene beakers (Plastikbecher GmbH, Giengen an der Brenz, Germany) filled with 150 mL10% N-medium and covered with perforated and transparent lids (four groups per beaker). Plants from the four different groups were marked with specific colors to follow the individual lineages; colors were randomized among treatments. To reduce algal growth, we changed the medium every two to three days and wrapped the sides of the beakers with three layers of black plastic foil to decrease light incidence. Inside the plastic beakers, we propagated each lineage for five generations using single descendant propagation. To assess plant fitness and resistance to copper excess, we moved the first and second daughter plant of generation five into separate 250 ml polypropylene beakers containing control conditions (10% N-medium) or recurrent copper excess (20 µM CuSO_4_ in 10% N-medium); the first and second offspring were randomly assigned to the treatment groups. After setting up the experiment, we took a picture for surface area analysis using a camera installation with a webcam (HD Pro Webcam C270, Logitech, Lausanne, Switzerland; webcam software 2.12.8). To reduce algae growth, we exchanged the medium after four days, and we took a second picture after eight days of growth.

### Statistical analyses

We calculated growth rates based on frond surface area [(End value)-(Initial value)]/days. Additionally, we estimated resistance to copper excess as the ratio in the surface area of plants grown under copper excess relative to the mean value of plants grown under control treatment). To test whether plant resistance and growth rates under control conditions and copper excess are affected by the selection regimen and the pre-treatment, we fitted a linear mixed effects model using selection regimen and pre-treatment as fixed effects, and as random effects the pair of a control and a copper pre-treatment ponds (“pair”) and blocks of ponds (groups of three to four ponds, “block”) regarding their physical position in the field (∼Selection*Pre-treatment+(1|pair)+(1|block)). Models were run in R v4.3.1 using the package lme4 v1.1.34 (Bates et al., 2015), and *P*-values of the fixed effects were estimated with Type III Analysis of Variance with the Satterthwaités method using anova of the package stats (R Core Team, 2023). We used the packages DHARMa v0.4.6 (Hartig, 2022) to verify whether the scaled residuals of our fitted models correlated to the simulated values from the same model, and effects v4.2.2 (Fox & Weisberg, 2018) to predict the effects of the fitted model. When analyzing plant resistance and fitness under copper excess, we found an influential sample in the copper pre-treated large population under copper environment, which showed Cook’s distance higher than 0.5, low correlation between the standardized residuals and theoretical quantiles (Q-Q plot) and whose corresponding sibling in the control environment was lost. Therefore, we excluded this outlier from the analysis. Plots were displayed with ggplot2 v3.4.3 (Wickham, 2016) and data was organized with readxl v1.4.2 (Wickham & Bryan, 2023), data.table v1.14.8 (Dowle & Srinivasan, 2023), tidytext v0.4.1 (Silge & Robinson, 2016) and dplyr v1.1.2 (Wickham et al., 2023). Figures were edited with Inkscape (Inkscape Project, 2018).

## RESULTS

### Proof-of-concept: duckweeds possibly adapt through both the stochastic and deterministic epigenetic routes

To assess whether *S. polyrhiza* acquires resistance to copper excess through either the deterministic or stochastic route, we measured plant resistance of single descendants and large populations grown under control and copper pre-treatment, the latter reduced plant growth rates outdoors by 16 % (Supporting Information Fig. S1). Both the presence of selection and copper pre-treatment affected plant resistance: regardless of the pre-treatment, plants from the large populations tended to have higher resistance to copper excess than plants from the single descendant lineages (Fig. 1b; selection: F_(1,3.33)_ = 7.24, *P* = 0.066, linear mixed effects model). Furthermore, pre-treatment in copper stress tended to increase copper resistance in the single descendants but not in the populations (Fig. 1b; selection x pre-treatment: F_(1,12.78)_ = 7.48, *P* = 0.017, linear mixed effects model).

To differentiate whether the increase in resistance was due to lower growth under control conditions or higher growth under copper excess, we analysed plant fitness under both treatment environments separately. Under control conditions, neither the population size nor the pre-treatment altered plant fitness (Fig. 2c; F_(1,10.33)_ = 0.50, *P* > 0.1, linear mixed effects models). In contrast, under copper excess, plants from the large populations outperformed plants from the single descendant lineages regardless of the pre-treatment (Fig 2d; selection: F_(1,3.68)_ = 8.34, *P* = 0.049, mixed effects models). Furthermore, pre-treatment in copper stress tended to increase plant fitness under copper excess in the single descendant lineages but not in the populations (selection x pre-treatment: F_(1,12.79)_ = 3.86, *P* = 0.071, linear mixed effects models). Therefore, variations in copper resistance were mainly due to variations in growth rates under copper excess and not under control conditions.

## DISCUSSION

Here we presented an experimental approach to test a long-standing controversy: whether epigenetic inheritance can lead to rapid adaptation to environmental stresses through stochastic or deterministic epigenetic variation. Our proof-of-concept experiments suggests that both the deterministic and stochastic routes may alter duckweed fitness under recurring stress.

### Proof-of-concept experiment

Our experiment suggests that duckweed populations may acquire stress resistance through the stochastic route, as, regardless of the pre-treatment, offspring from large populations outperformed offspring from the single descendants. The higher fitness of the large populations compared to single descendants may have been caused by two factors: first, variations among single descendants and large populations may have been confounded by our method to identify the single descendants during the pre-treatment phase. To identify the single descendants, we marked the single descendants throughout the pre-treatment phase. Although we also marked plants from the large populations in the last two generations of the pre-treatment, we cannot fully rule out that the prolonged marking of the single descendants affected plant fitness even after the five generations of growth under shared control conditions prior fitness assay. In this case, the difference between the large populations and single descendants is not caused by selection (stochastic route) but by the marking, a pre-treatment effect (deterministic route) that would have persisted across five generations. Second, the higher fitness of the large populations than of the single descendants may have been caused by selection of stochastic variants. Our experiment was limited to five generations. While such short durations are sufficient for natural populations to become adapted through selection of standing genetic variations (Züst et al., 2012), it is unclear whether monoclonal populations may acquire sufficient epigenetic variation for such rapid adaptation taking place. Future experiments that assess variation in plant fitness among single descendant lineages, and long-term experiments that ensure that small and large populations experience the very same environment during the pre-treatment phase will be needed to differentiate among these two possibilities.

While the largest differences in fitness was caused by the different population size, we also obtained evidence for deterministic effects (interaction of population size and pre-treatment on plant resistance): copper pre-treatment increased fitness of stress single descendants and decreased fitness of the population plants. This is in line with recent studies suggesting that the previous environment alters plant fitness and epigenetic marks across multiple clonal generations (Anastasiadi et al., 2021; Benson et al., 2021; Colinas et al., 2023; González et al., 2018; Li et al., 2018; Sammarco et al., 2023). Future manipulative studies are needed to test for specific mechanisms of the deterministic route to confirm the presence of this phenomenon and to understand whether the observed routes are truly caused by epigenetic inheritance *sensu stricto* (genome-associated mechanisms) or by other non-genetic inheritance mechanisms.

While the proposed experiment is not designed to tease apart the specific epigenetic mechanism, the approach can – if carried out carefully – resolve the contribution of the deterministic and stochastic route in a wide range of organisms. In addition, the approach can also be used to assess the burdens of genetic mutation accumulation, as proposed previously (Ho et al., 2021; Ho et al., 2020). We would like to highlight a few points to make maximal use of this experimental approach:

### Inferring the stochastic and deterministic route

The above-described experiment with duckweeds reveals a critical issue: the pre-treatment environment of the small and large population must be equal, otherwise one cannot differentiate between the stochastic and deterministic route. In our experiment, this could be achieved by caging small populations inside the large population – this would alleviate the potential bias of the marking. However, one must ensure that the small populations still experience the same environment as the large populations despite the caging. In duckweeds, this can be achieved by separating the small populations from the large populations using floating disks while keeping the density of the small and large populations equal. It is of utmost importance to carefully design of the experiments considering the specific biology of each organism.

So far, we treated the stochastic and deterministic route as separate, non-overlapping entities. This may be an unrealistic assumption. For instance, deterministic epialleles may not be induced in an entirely deterministic manner; environmental stresses may simply increase the likelihood of an epiallele to appear but not cause all individuals changing their epigenetic status. Selection could thus still act on environment-induced “deterministic” epialleles. Similar accounts for the stochastic epimutations: environmental stresses may increase the likelihood that certain epimutations appear, associated to the presence of epigenetic hotspots (Hazarika et al., 2022; Zheng et al., 2017), and thus epimutations may not be entirely stochastic. Are the stochastic and deterministic route therefore two extremes of the same process? One argument holds against this: in *A. thaliana*, spontaneous epimutations mostly accumulate in the CG context (Becker et al., 2011; Denkena et al., 2021; Schmitz et al., 2011; van der Graaf et al., 2015) whereas environment-induced epialleles are most prominent in the CHG and/or CHH context (Annacondia et al., 2021; Lin et al., 2022; Wibowo et al., 2016; Yadav et al., 2022; Zhou et al., 2019). This supports the idea that the stochastic and deterministic route are, at least in the context of DNA methylation in plants, two fundamentally different processes.

### Inferring the molecular mechanisms

Another critical issue is the interpretation of the results: is the underlying molecular mechanism of genetic or epigenetic nature? Even when using highly inbred or clonally reproducing species, genetic mutations may accumulate and account for variation in phenotypes or fitness, particularly in the large populations. We suggest following experiments and analyses to tease apart genetic from epigenetic effects: first, unbiased methods (e.g. high-throughput sequencing) should be deployed to screen for epigenetic variations. Second, transcriptome and proteome data should be generated to link fitness to phenotypes to gene expression and epigenetic variation. Third, differentially regulated genes or epigenetic machineries should be manipulated both genetically as well as chemically to test whether the identified genes or epigenetic machinery affect stress resistance or the process of stress adaptation. Fourth, phenotypes or fitness should be continuously assessed for multiple generations after stress release – if variations between groups have an epigenetic basis, the variation should diminish or at least change in some lineages over time. Fifth, high-throughput sequencing should be deployed to screen for genetic variation in identified genomic regions. While in isolation none of these approaches will provide a clear answer whether genetic or epigenetic mechanisms are at play, combining these approaches will be powerful to disentangle genetic and epigenetic factors.

## CONCLUSIONS

Taken together, we here provide an experimental framework to assess the relative contribution of deterministic epialleles and selection of stochastic epigenetic variants to rapid stress adaptation. Through this approach, carefully designed experiments can provide novel insights whether and how epigenetic inheritance may lead to rapid stress adaptation. Considering the on-going loss of intra-specific genetic diversity and the rapid pace of environmental change, these insights may become increasingly relevant when assessing species potential to resist global change.

## ACKNOWLEDGEMENTS

We would like to thank Pauline Prüsener, Anke Grosse-Brinkhaus, Maite Görtz Lizarraga and the Botanical Garden of the University of Münster for supporting the outdoor experiment and Shuqing Xu for providing facilities. This study profited from interactions within the GenEvo Research Training Group 407023052/GRK2526/1 funded by the German Research Foundation (DFG). The work was funded by the Volkswagen Foundation (Grant Nr 97236) to MH and the German Research Foundation to MH (grant Nr. 512079118). The project was supported by the University of Münster and the University of Mainz.

## AUTHOR CONTRIBUTION

MH conceived the ideas and designed methodology, AC collected the data, AC and MH analysed the data, MH and AC drafted the manuscript. Both authors contributed critically to the drafts and gave final approval for publication.

## STATEMENT ON INCLUSION

This study was discussed and carried out by a group of scientists originating from different continents and by technicians from the local region. All authors were engaged in the design of the experiments to ensure that a diverse set of perspectives were included. Efforts were made to consider relevant work published in the local language.

## DATA AVAILABILITY

The raw data of the outdoor experimental evolution experiment will be archived on Dryad upon acceptance.

