## Supporting Information Figure S1 for "Assessing rapid adaptation through epigenetic inheritance: a new experimental approach"

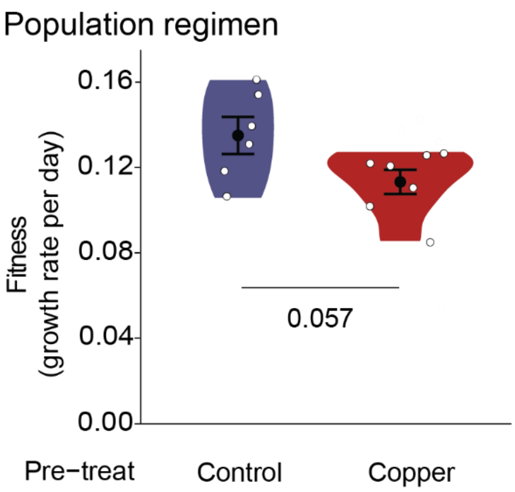


***Figure S1. Plant fitness of the population regimen when grown under control and copper excess.*** *Fitness values were measured after 12 days of free growth.* P-*values refer to linear mixed effects models. The black circles mark the means, and error bars show the standard errors. N = 6-7. Pre-treat = Pre-treatment.*
